# Density dependence promotes species coexistence and provides a unifying explanation for distinct productivity-diversity relationships

**DOI:** 10.1101/2025.07.28.667229

**Authors:** Liang Xu, Christopher A. Klausmeier, Emily Zakem

## Abstract

Understanding diversity patterns in complex communities, such as microbial consortia, requires a mechanistic framework appropriate for many species. Negative density dependence is often utilized in complex ecosystem models, typically as a density-dependent mortality term for a population, but its full impact on community structure remains unclear. Here we use mechanistic population models of resource consumption to examine the effects of negative density dependence and develop a tractable framework for understanding diversity patterns in complex systems. To provide mechanistic grounding, we quantify how density-dependent mortality expands coexistence zones along resource gradients in simple communities using graphical analysis. We then derive an analytical, ecologically insightful formula predicting species abundances in subsets (guilds) of complex communities, in which many species share a resource or predator. Finally, we use the formula to explain how distinct relationships between productivity and diversity emerge from the resulting mechanistic framework, providing insights into previously unreconciled observed patterns.

## 1. Introduction

Since Hutchinson articulated the “paradox of the plankton” (Hutchinson 1961), numerous hypotheses have been proposed to explain the coexistence of many plankton species competing for a limited number of resources. Resource competition theory (RCT), a foundational framework in ecology, has deepened our understanding of plankton diversity (Tilman 1982; Litchman & Klausmeier 2001; Klausmeier *et al*. 2004; Litchman & Klausmeier 2008). However, RCT analysis is typically restricted to simplified food web structures (Holt 1977; Holt *et al*. 1994; Chase & Leibold 2003), where its graphical techniques are not easily extended to more than a few (< 3) resources and species. Microbial communities, which contain thousands of species (or species-like equivalents) consuming diverse substrates (Moran *et al*. 2022), therefore present a significant challenge to the application of this theoretical framework. To understand species coexistence and diversity patterns in such complex communities, we need a general, comprehensive, and mechanistic framework applicable to diverse systems with many resources and species.

Any such framework must consider the impact of negative density dependence, the suppression of a species’ net growth rate as its density increases due to multiple potential biological processes, such as viral lysis (Winter *et al*. 2010; Thingstad 2022), host-specific parasitism (McPeek 1998), and soil pathogen interactions (Bagchi *et al*. 2014; Comita *et al*. 2014; LaManna *et al*. 2017a; Adler *et al*. 2018; Tuck *et al*. 2018). Though phenomenological, density-dependent mortality in mathematical descriptions of population growth captures processes that are too uncertain or complex to model explicitly, and so is often utilized in studies of complex ecosystems (Bagchi *et al*. 2014; Fricke & Wright 2017; Adler *et al*. 2018; Bendik & Dries 2018; Xu *et al*. 2022). Additionally, density-dependent mortality is a well-established mechanism for resolving the paradox of the plankton because it facilitates the persistence of inferior competitors (Chesson 2000; Thingstad 2000).

Although density-dependent mortality is widely invoked to explain biodiversity patterns (Holt & Lawton 1993; Winter *et al*. 2010; Bagchi *et al*. 2014; Comita *et al*. 2014; LaManna *et al*. 2017b), theoretical studies, grounded in mechanistic resource competition models, remain limited relative to empirical studies. Thus, a key question remains: how does density dependence shape species coexistence and relative abundance patterns along resource gradients? Although theory has shown that density dependence can stabilize coexistence and allow more species to persist than there are limiting resources (Leibold 1998; Adler *et al*. 2007; McPeek 2012, 2019), most analyses rely on low-dimensional isocline approaches, leaving its role in more diverse systems poorly understood.

A broader question is how biodiversity responds to productivity and how this relationship varies. Different productivity–diversity relationships (PDRs) have been observed, with monotonic increases and unimodal (hump-shaped) curves common (Mittelbach *et al*. 2001). For example, Fraser et al. (2015), analyzing 30 grassland sites on six continents, found strong support for a unimodal PDR, with richness peaking at intermediate productivity at regional and global scales. However, Adler et al. (2011) sampled 48 herbaceous plant communities across five continents and found no consistent relationship: while some individual sites showed linear or hump-shaped trends, the overall pattern lacked coherence.

These observed PDRs are not easily explained by classic resource competition theory alone, since the number of distinct resources sets a limit on the number of stably coexisting species, and it is unlikely that resource diversity increases along productivity gradients at the same pace as species diversity. Additional mechanisms are therefore required. Proposed mechanistic explanations include spatial or temporal heterogeneity (e.g., due to dispersal or time-varying resource supply) (Abrams 1988; Leibold 1996), sampling limitations (Pastor *et al*. 1996), and spatial scale dependency (Steiner & Leibold 2004), but any one of these mechanisms cannot explain why multiple distinct patterns may arise. Density-dependent mortality may constitute another potential mechanistic explanation, but this has not yet been rigorously evaluated. Exploring this potential requires a theoretical framework that can mechanistically link PDRs to trophic interactions and ecosystem structure that is applicable to a large number of species. Ideally, such a framework would also explain how different PDRs may emerge from a general description of ecosystem dynamics.

Here, we develop a unified framework that links density dependence to both species coexistence and PDRs. We begin by extending classic resource competition theory to reveal how density dependence modifies coexistence criteria, reshaping resource consumption vectors and supply points. We use well-established, simple food web modules (Holt 1997) to provide this mechanistic grounding, illustrating how density dependence enables more species to coexist on limited resources (Section 2). Building on this foundation, we then derive general formulas for species abundances within analytically tractable subsets of a complex community: a bottom-up (one shared resource) or a top-down (one shared predator) guild (Section 3). These formulas are applicable to systems with many species, such as diverse microbial consortia with one dominant predator. Finally, we demonstrate how this framework, combined with trait trade-offs, captures multiple PDRs in consistency with observations, which unifies coexistence theory and PDRs within one mechanistic description (Section 4).

## 2. Graphical analysis of simple systems

We begin by exploring how density-dependent mortality affects species coexistence in simple systems using graphical analysis, building on previous work (Leibold 1996; Chase & Leibold 2003). Specifically, we quantify how density-dependent mortality expands the coexistence domain in three well-known food web modules: (1) two species competing for two substitutable resources, (2) three generalists competing for two resources, and (3) a diamond food web with two prey species and one shared predator.

### 2.1 Two species and two resources

We first consider a food web of two species competing for two nutritionally substitutable resources. Assuming linear functional responses, species *i* with density *N*_*i*_ grows at rate *μ*_*ij*_ on resources *R*_*ii*_, supplied with concentration *s*_*j*_ and dilution rate *a*:

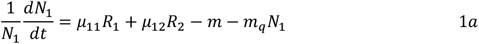

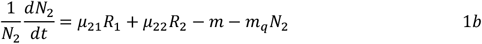

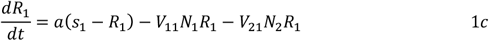

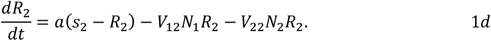

Here *V*_*ij*_ are consumption rates with yields *Y*_*ij*_ = *μ*_*ij*_/*V*_*ij*_ . Density-dependent mortality for species *i* with density *N*_*i*_ is modeled as 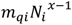, where *x* > 1 . For simplicity, we use quadratic mortality (*x* = 2) in the main text. We generally assume equal linear mortality rates (*m*) and quadratic mortality constants (*m*_*q*_). Species-specific cases can be rescaled to the general framework, providing quantitative but not qualitative variability to results (Supplementary Material).

Without density-dependent mortality (*m*_*q*_ = 0), coexistence requires each species to be competitively superior on one limiting resource, represented by intersecting ZNGIs (zero net growth isoclines) (Ryabov & Blasius 2011; McPeek 2014; Kleinhesselink & Adler 2015; Letten *et al*. 2017) (Fig. 1b). Density-dependent mortality does not alter invasion criteria—resource levels must still lie above a species’ invasion ZNGI where *N*_*i*_ = 0—but shifts equilibrium resource levels 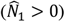 to depend on species densities and thus on supply rates. Equilibrium ZNGIs are:

**Figure 1:**
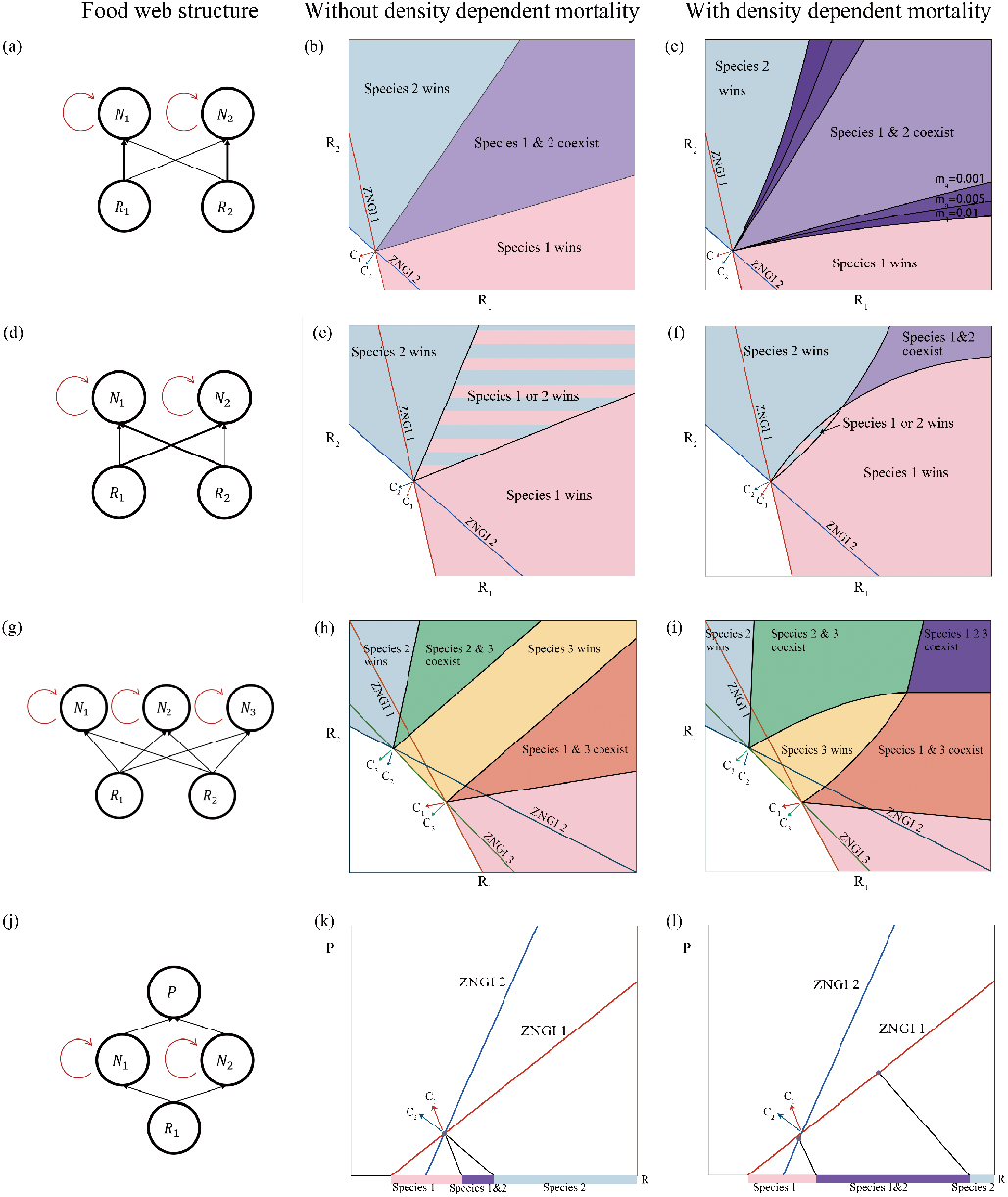
Illustration of the range of resource supply rates for the species coexistence with and without the density-dependent effect. The impact vectors of species on resources are denoted by *c*_1_, *c*_2_, *c*_3_. (a), (d), (g), (j). The food web structures. (b). The two generalists can coexist when *m*_*q*_ = 0; (d). Increasing *m*_*q*_ expands the range of resource supply rates for species coexistence. The model parameters are *μ*_11_ = 1.0, *μ*_21_ = 0.2, *μ*_12_ = 0.5, *μ*_22_ = 0.5, *V*_11_ = 3.3, *V*_21_ = 0.67, *V*_12_ = 1.67, *V*_22_ = 1.67, *m* = 0.1, *a* = 1; (e). The two generalists cannot coexist when *m*_*q*_ = 0 because their impact vectors indicate a priority effect; (f). The effect of density dependence facilitates coexistence at higher supply rates. For the supply rates vector falls in the striped zone, the priority effect emerges. The model parameters are *μ*_11_ = 1.0, *μ*_21_ = 0.2, *μ*_12_ = 0.5, *μ*_22_ = 0.5, *V*_11_ = 1.1, *V*_21_ = 1.8, *V*_12_ = 2.2, *V*_22_ = 0.6, *m* = 0.1, *a* = 1. (h). When *m*_*q*_ = 0, at most two species can coexist in the supply rate space. The outcome of species interaction depends on where the vector of resource productivities falls.; (i). When *m*_*q*_ = 0.01, the two pairwise coexistence zones overlap, indicating an area where three species can coexist. The model parameters are a = 1, *μ*_11_ = 0.27, *μ*_22_ = 0.24, *μ*_13_ = 0.15, *μ*_23_ = 0.15, *V*_11_ = 0.9, *V*_22_ = 0.8, *V*_12_ = 0.5, *V*_22_ = 0.5, *m* = 0.1, *a* = 1. (k-l). In the diamond food web, two species can coexist at a larger range of supply rates with increase of *m*_*q*_; (k).*m*_*q*_ = 0; (l). *m*_*q*_ = 0.01; The boundaries are given by Eqns. 4 and 5. The model parameters are *μ*_1_ = 2.8, *μ*_2_ = 1.2, *V*_1_ = 4.2, *V*_2_ = 1.8, *g*_1_ = 8.7, *g*_2_ = 1.725, *f*_1_ = 5.8, *f*_2_ = 1.15, *m* = 0.1, *m*_*p*_ = 0.1, *a* = 1.

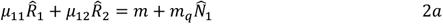

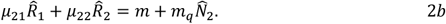

where equilibrium conditions are denoted by 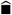’s. Unlike invasion ZNGIs, these equilibrium ZNGIs with 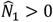 vary with biomass and elevate equilibrium resource levels (Fig. S2). As supply increases, mortality rises with density, raising resources. Strong competitors benefit from increased growth but also suffer higher mortality, while weaker ones, at lower densities, experience less mortality. This asymmetry broadens the coexistence domain beyond what is possible under pure resource competition (Leibold 1998; McPeek 2019).

We next quantitatively analyze the impact of density-dependent mortality on the boundaries between competitive exclusion and coexistence regions using mutual invasibility analysis (Chesson 2000) (Fig. 1c) (Supplementary Material). Density-dependent mortality significantly widens coexistence ranges and produces nonlinear boundaries between competitive exclusion and coexistence regions (Fig. 1c). The lower boundary of the coexistence zone curves downward compared to the curve of *m*_*q*_ = 0. This indicates that species 2 can invade at a lower supply rate of resource 2 when species 1 dominates, since species 1 cannot deplete resources to its invasion ZNGI (*μ*_11_*R*_1_ + *μ*_12_*R*_2_ = *m*) but only to the higher equilibrium 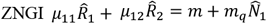. The same principle applies to the upper boundary of the coexistence zone when species 1 can invade a community with the resident species 2.

In general, a larger *m*_*q*_ results in a larger coexistence zone, indicating that stronger density-dependent mortality promotes species coexistence across an expanded range of resource supply rates (Fig. 1c). Box 1 summarizes the four possible competitive outcomes.

#### Density-dependent mortality reduces the likelihood of priority effects

Notably, density-dependent mortality allows coexistence where two generalists would otherwise exhibit priority effects (Chesson 2000; Fukami 2015; Grainger *et al*. 2019). In Fig. 1d-f, we selected consumption rates for the two species where each species consumes more of its less preferred resource 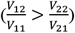, normally yielding first-come-first-survive without density dependent mortality. The striped area on the resource plane in Fig. 1e represents where priority effects occur: the species that initially establishes itself in the community excludes the other. Without density-dependent mortality, this region expands linearly with resource supply, with no intersection of the boundaries (Fig. 1e). In contrast, with density-dependent mortality, the invasion boundaries intersect at sufficiently high resource supply levels (Fig. 1f). Thus, low supply maintains the priority effect, but high supply enables coexistence. This finding suggests that both supply magnitude and ratio govern the ecological outcomes.

### 2.2 Three species and two resources

We next examine how density-dependent mortality can expand not only the domain of coexistence but also increase the total number of coexisting species. We consider a food web with three species and two substitutable resources, with two generalists (species 1 and 2) with advantages on different resources, and one generalist (species 3) having intermediate advantages on both resources. Without density-dependent mortality, the system exhibits two distinct regions of pairwise coexistence. As in the above example, density-dependent mortality expands the coexistence regions by bending the invasion boundaries (Fig. 1g-i). In this case, the two coexistence zones may overlap at high resource supply rates, leading to coexistence of all three species (Fig. 1i).

### 2.3 The diamond food web: Two species, one predator, and one resource

We finally demonstrate how density-dependent mortality promotes species coexistence in a different classic system: the “diamond” food web with two species *N*_*i*_ competing for one resource *R* and consumed by a common predator *P* (apparent competition (Holt 1977))as:

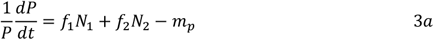

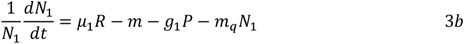

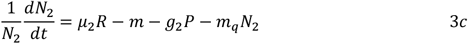

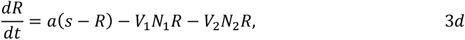

where *f*_*i*_ is predator growth at predation rate *g*_*i*_ on species *i*, and *m*_*p*_ denotes the mortality rate of the predator. Without density-dependent mortality, coexistence requires the two species’ ZNGIs to intersect on the predator-resource plane (Fig. 1k). The stability of this equilibrium depends on the condition that the species with superior resource-competitive capabilities is more poorly defended against the predator, while the worse resource competitor is better-defended against the predator (Leibold 1996). Here, we assume that species 1 is the better resource competitor and species 2 is better defended against the predator, i.e., *μ*_1_ > *μ*_2_, *g*_1_ > *g*_2_.

With density-dependent mortality, the two critical resource supply concentrations are given by:

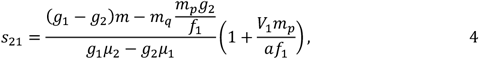

And

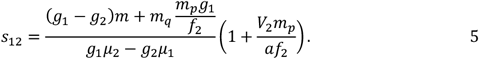

where *s*_*ij*_ denotes the threshold of the supply rate for species *i* invading species *jj* (Supplementary Material). Increasing *m*_*q*_ increases the interval between *s*_21_ and *s*_12_, promoting coexistence. We note that the invasion point of species 2 is still bounded by a minimum supply, which is determined by the intercept of its ZNGI and the supply axis. Unlike the previous cases (Fig. 1a-i), further increasing resource supply does not necessarily result in coexistence, as it ultimately goes beyond the coexistence interval, favoring the superior apparent competitor, species 2 (Fig. 1l). Thus, although density dependence continues to promote species coexistence, its effect is weakened by apparent competition as resource supply increases. The community composition then shifts towards species with higher predator resistance.

Note that the case of no density-dependent mortality (*m*_*q*_ = 0) is a special case of these general equations (Supplementary Material). The four possible competitive outcomes are summarized in Box 1 (Fig. 1i-l).

## 3. Control formulas for complex systems

We have quantified how density-dependent effects promote species coexistence, allowing more species to coexist than the number of limiting factors in simple systems. However, graphical analysis becomes impractical for quantitatively extending this approach to communities with more than a few species. We need a more general quantitative approach to mechanistically understand how diversity changes with resource productivity across different food-web structures in more complex communities.

To address this, we consider analytically tractable subsets of communities with many coexisting species. Specifically, we consider two types of subsets, which we refer to as guilds: (1) a bottom-up guild, in which multiple species compete for a single shared resource, while their interactions with higher-level predators remain unspecified (Fig. 2a); and (2) a top-down guild, in which multiple species are preyed upon by a single shared predator, with unspecified details of their resource use (Fig. 2b). This approach aligns with a common and ecologically realistic strategy of analyzing food-web modules—simplified sub-networks where species interactions are governed by basic trophic relationships, such as resource consumption and predation (Holt 1997)—while still allowing considerable complexity in the rest of the food web. For example, tropical forests, which harbor high plant diversity competing for similar resources (Gentry 1988), and marine phytoplankton, which compete for a limited set of inorganic nutrients (Hutchinson 1961), both exemplify bottom-up guilds. Top-down control is evident in freshwater lakes, where invertebrate and vertebrate predators structure distinct prey assemblages (Werner & McPeek 1994). Similarly, microorganisms of comparable size often share predators due to size-based predation (Clauset & Erwin 2008; Ward *et al*. 2012; Follett *et al*. 2022). Considering these guilds, each with a single shared limiting factor, permits analytical treatment that would be intractable in fully specified complex networks.

**Figure 2:**
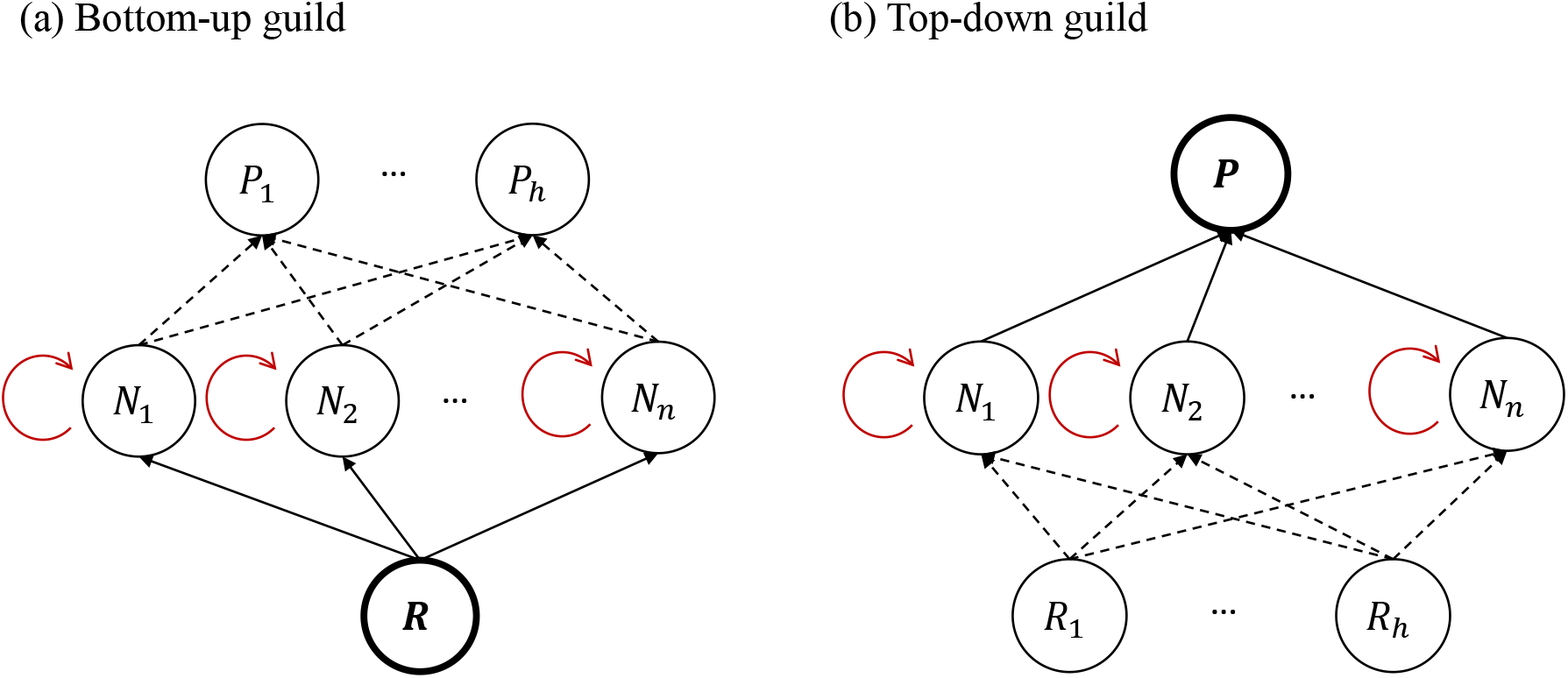
Two types of guilds illustrating species interactions within and between trophic levels. The negative density dependence is denoted by the red arrows. (a). The bottom-up guild assumes that consumer species are mainly limited by food availability of one common resource; (b) The top-down guild assumes that consumer species are restricted by the common predator. Predators are denoted by P; consumer species are denoted by N; resources are denoted by R.

We next develop analytical expressions to characterize coexistence patterns of these guilds, which allows us to later explore mechanistic relationships between productivity and diversity. We present a general control formula that results from our derivations of the two guild-specific control formulas. This general form provides ecological insight into coexistence dynamics, including the impact of density-dependent mortality (Section 3.1). We then present the specifics of the bottom-up formula (Section 3.2), noting that the corresponding specifics of the top-down formula follow a similar derivation and structure (Box 2). We use the bottom-up formula to link coexistence and productivity-diversity patterns within one framework (Section 3.3).

### 3.1 The general control formula

We derive control formulas for each of the guilds, which results in one general control formula for species abundances (see Box 2 for derivation overview and SM section 1). The resulting general formula describes the steady-state density 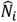of species *i* as:

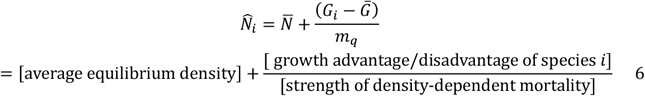

The quantity 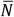 is the appropriately weighted average density, set by the predator or resource (details below). Thus, 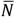is set by the species of the guild that can coexist (i.e., have a positive steady-state density) at equilibrium, which may be the complete set or a subset of all possible species. *G*_*i*_ is the net growth rate of species *i*, which we define as the net growth excluding the density-dependent term (i.e., *G*_*i*_ = ∑_*h*_ *μ*_*ih*_*R*_*h*_ − *m* − ∑_*h*_ *g*_*hi*_*P*_*h*_). Finally, 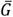 is the weighted average growth rate of sustained species in the community, with a specific form determined by the guild type.

Equation 6 thus explains the equilibrium density of a species as determined by three interdependent components. The first component, the average density of the community 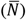, is intrinsically linked to the shared limiting factor (resource or predator) defined by the guild type. The second component is the discrepancy between the net growth rate (*G*_*i*_) of the focal species and the weighted community average. When the value of the focal species’ *G*_*i*_ surpasses that of the community average 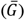, it represents a relative competitive advantage that elevates the species’ abundance above the average. The third component is the strength of density-dependent mortality 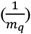, which scales the impact of the second component. When density-dependent mortality is negligible (*m*_*q*_ = 0), all coexisting species must have the same net growth rate 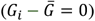. In contrast, when density dependence is strong, species with net growth rates lower than the community average may persist, increasing diversity of the guild at equilibrium.

### 3.2 The specific control formula for a bottom-up guild

In a bottom-up guild, multiple consumer species compete for and are predominantly limited by a single shared resource. Predation is unspecified and thus not the main constraint on total consumer density (Fig.3a). The system is described as:

**Figure 3:**
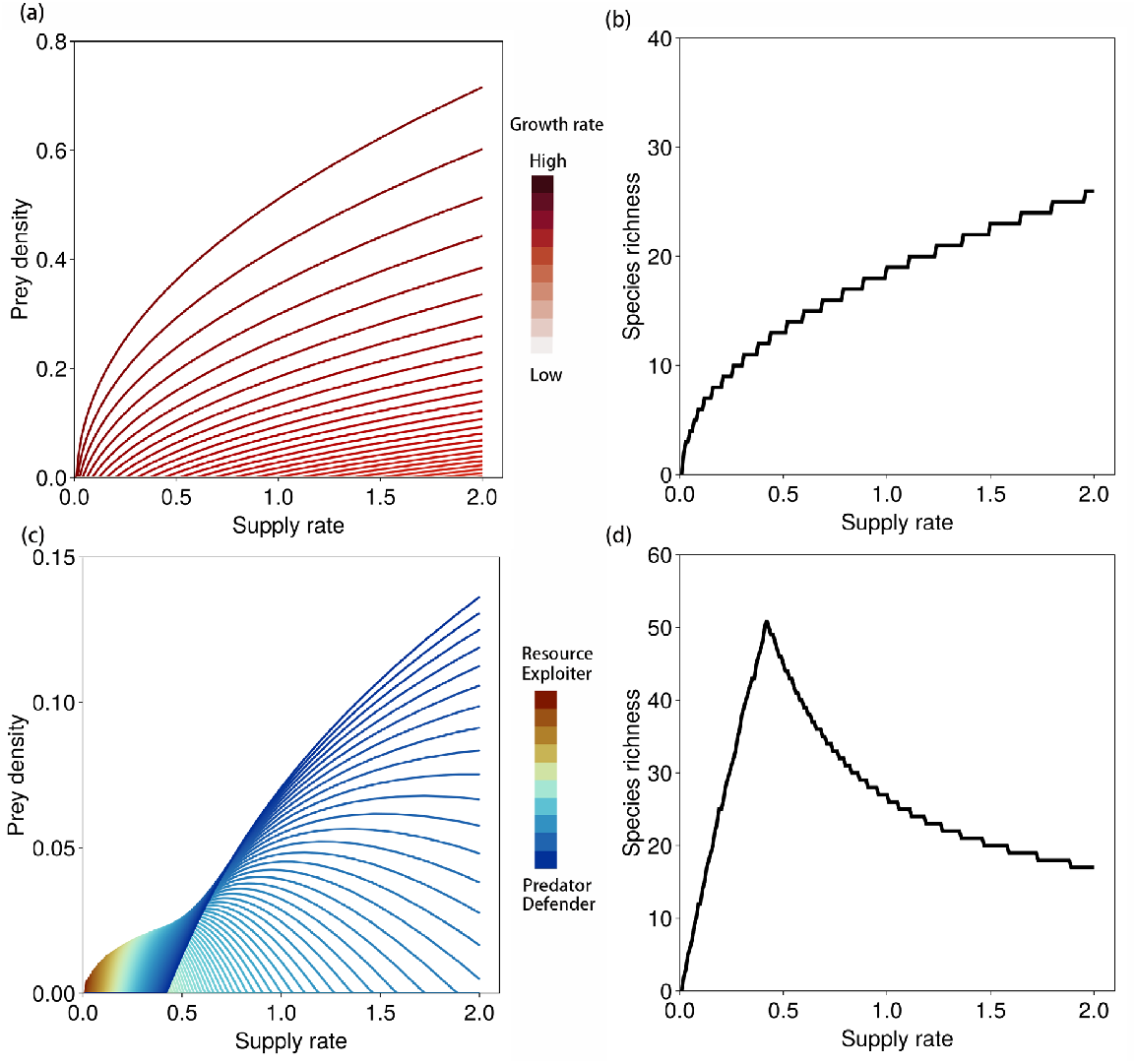
Negative density dependence generates contrasting productivity–diversity relationships. (a) Simulation of a bottom-up guild with 100 species ranked by growth rates (color gradient). (b) In the presence of density-dependent mortality, species richness increases with resource supply. (c) Simulation of a diamond food web with 100 species, sharing a common predator and resource. Species are ranked by growth and predation rates. (d) Under density-dependent mortality, the resulting productivity–diversity relationship becomes unimodal. As in Section 2, species are ordered by decreasing intrinsicgrowth rates (*μ* > *μ* > ⋯ > *μ*) for simulations in (a) (b) and with additional condition 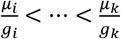 for simulations in (c) and (d).

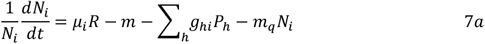

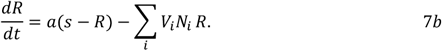

with parameters as defined above (Eqn. 1). The specific form of the control formula (Eq. 6) for the bottom-up guild is

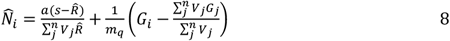

where the average density is 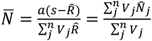, representing the resource consumed 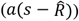 divided by the total consumption rate of all sustained species 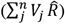. Though predation is not specified, predation does indirectly impact the effective resource level 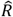, because predation reduces prey densities and thereby indirectly influences the equilibrium resource concentration 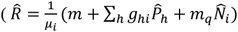 . The average community growth rate is weighted by each species’ resource consumption rate 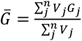, where *G*_*j*_ = *μ*_*j*_ *R* −*m* − ∑_*h*_ *g*_*hk*_ *P*_*h*_. Thus, Eqn. 8 reveals a trade-off between species density and diversity (which also applies to the top-down guild). As the strength of density-dependent mortality (*m*_*q*_) increases, more species can coexist, but the average density per species declines, as made clear by the simpler derivation noted in Box 2.

### 3.3 Linking coexistence and productivity-diversity relationships

The control formula, using the specifics for the bottom-up guild, links resource supply, traits, and species abundances (and thus diversity) mechanistically, allowing prediction of diversity patterns as a function of resource supply. For example, Eqn. 8 can be used to determine the minimum resource supply rate required for a species to persist, as follows. First, we note that the equilibrium resource concentration is associated with the resource supply rate *s* (Eqn. 7b and 15) as:

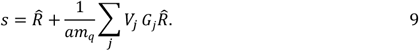

given an existing set of species (1 to *jj*). We next consider how species *i* may invade relative to this rate. Imposing 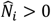 in Eqn. 8, the threshold supply rate for species *i* corresponds to the condition under which its equilibrium density becomes positive:

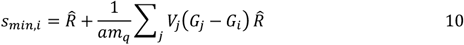

This expression defines the minimum supply rate necessary for species *i* to persist, given the presence of other coexisting species. The summation term captures the cumulative effect of fitness differences weighted by resource uptake rates between species *i* and other species that coexist at lower supply rates. Note that 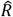 is the same in Eqn. 9 and 10 when species *i* is just invading 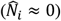, and thus that 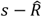quantifies the amount of resource consumed by the community. If *s* > *s*_*min,i*_, the consumed resource exceeds the minimum requirement, allowing species *i* to invade and coexist with the established community.

Furthermore, we note that 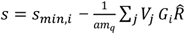. Thus, when *G*_*i*_ remains positive across the supply gradient, the coexistence criterion is always satisfied. Conversely, if *G*_*i*_ becomes negative, the criterion is violated. This yields a tractable condition for quantifying coexistence and diversity for any specified set of species. Therefore, by examining how species’ net growth rates vary with resource supply, we can directly link resource-driven productivity to biodiversity patterns, as we next demonstrate.

## 4. Mechanistic explanations for distinct productivity-diversity relationships

Our mechanistic framework links resource supply rate, which governs the productivity of the ecosystem, to species richness and diversity. Here, we explore how this provides a mechanistic basis for the two most commonly observed PDRs: increasing and unimodal. We identify the factors that determine whether one or the other should emerge and use our results to interpret observations in new ways.

### 4.1 Increasing productivity-diversity relationship

We continue to use the control formula, specific for a bottom-up guild, to show how diversity increases with productivity via the resource supply rate. Critically, this makes the assumption that predation has negligible impact on species coexistence patterns. This may, for example, describe phytoplankton communities competing for scarce nutrients with little grazing pressure in oligotrophic systems (Reynolds 1984).

We illustrate the ecological dynamics underlying the increasing PDR clearly by using a specific example of this guild structure with simple tradeoffs between species traits. In a purely bottom-up guild with negligible predation, the net growth rate is *G*_*i*_ = *μ*_*i*_*R* − *m*, which is a monotonically increasing function of the equilibrium resource concentration 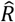. With density-dependent mortality (*m*_*q*_ > 0), the equilibrium resource concentration increases with the supply rate (Supplementary Material, Fig.S1, S3):

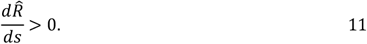

which in turn gives

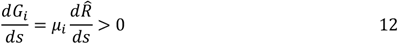

Thus, *G*_*i*_ increases with the supply rate, and the coexistence condition 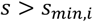 always holds once species *i* has successfully invaded. This mechanism produces an increasing PDR (Fig. 3b).

When density-dependent mortality is negligible (*m*_*q*_ = 0), the denominator diverges in Eqn. 10,and any differences in the net growth rates *G* are effectively amplified, implying that no finite supply rate can support the persistence of an inferior competitor. From the perspective of resource competition, the species with the lowest *R\** of the community can deplete the resource to its *R*^∗^, which does not increase with the supply rate. Thus, it prevents the other species from invading.

More generally, this framework indicates that an increasing PDR is expected whenever the net growth rate *G* remains stable or increases with productivity for the majority of species. This outcome is fundamentally shaped by density-dependent mortality, which acts as the key regulatory mechanism enabling the pattern. Importantly, this outcome is not restricted to a purely bottom-up guild without predators. Even when predation is present, increasing PDRs can still emerge provided that apparent competition is weak, such as if predation is sufficiently specialized to avoid driving species to extinction through shared predators (Holt 1977). Thus, increasing PDRs arise not only in pure bottom-up systems but also in predator–prey systems with specialized or low rates of predation (Fig. 4a, Supplementary Material section 3.5). We next explore how apparent competition via predation qualitatively changes the shape of PDR.

**Figure 4:**
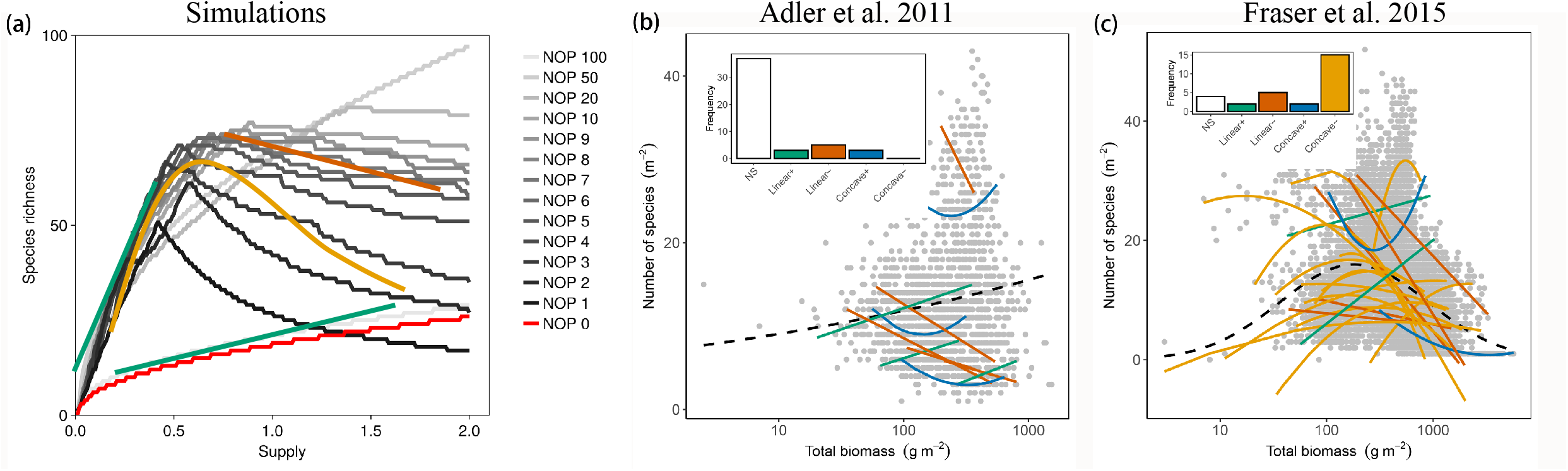
Density-dependent mortality generates productivity–diversity relationships (PDRs). (a). Simulated PDRs in a food web with 100 prey species and varying number of predators (NOP=0,…, 100). Different PDR shapes emerge at different ranges of resource supply rates (indicated by colored lines). Solid green lines: increasing PDRs; solid yellow lines: unimodal PDRs; solid brown lines: decreasing PDRs; solid blue lines: concave up PDRs. (b); (c). Empirical PDRs compiled from Adler et al. 2011 and Fraser et al. 2015 (see Supplementary Material). Solid lines show within-site regressions; dashed lines indicate regressions pooled across sites. Inset panels show the distribution of observed PDR types: NS = non-significant, Linear+ = positive linear, Linear− = negative linear, Concave+ = anti-unimodal, and Concave− = unimodal relationships.

### 4.2 Unimodal productivity-diversity relationship

We next illustrate how a qualitatively different, unimodal PDR emerges from the same general description of ecosystem dynamics, but when the details regarding predation are different. Specifically, we add a shared predator to the above bottom-up guild, resulting in the diamond food web structure, the simple but ecologically relevant case where prey species share a single resource and a common predator (explored using graphical analysis in Section 2.3) (Levin 1970; Holt *et al*. 1994; Leibold 1996). Thus, in contrast to the bottom-up guild, apparent competition from shared predation is significant. As in Section 2.3, we assume a trade-off between intrinsic growth rate and vulnerability to predation, reflecting a range of strategies from efficient resource exploiters (high *μ*, high *g*) to predator-resistant species (low *μ*, low *g*).

In this system, the net growth rate *G*_*i*_ = *μ*_*i*_*R* − *m* − *g*_*i*_*P* is no longer monotonic with resource supply rate. The derivative of *G*_*i*_ with respect to *s* takes the form

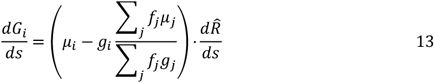

indicating that *G* exhibits variable responses to changes in the supply rate *s* and species diversity (Supplementary Material).

As above, increasing the supply rate increases the resource concentration, allowing more species to invade. However, the term 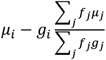 decreases when more species that are more resistant to the predator (with sequentially smaller *μ* and *g* than the *i*^*th*^ species) invade (Supplementary Material). Consequently, as the resource supply rate continues to increase, more predation-resistant species are established, reducing *G*_*i*_ . Eventually, *G*_*i*_ becomes negative, leading to the exclusion of fast-growing but predator-vulnerable species, and the coexistence criterion *s* > *s*_*min,i*_ is no longer satisfied. This process unfolds sequentially along the supply gradient.

Meanwhile, a shared predator can only persist once total prey biomass exceeds a threshold, ∑_*i*_ *f*_*i*_*N*_*i*_ > *m*_*p*_ (Eqn. 16 in Box 2). Our framework therefore indicates that a sufficient resource supply is needed to enable predator persistence, leading to apparent competition, which weakens density dependence among prey. As a result, diversity peaks at intermediate productivity, giving rise to a unimodal productivity–diversity relationship.

### 4.3 The emergence of multiple PDRs within one framework

Thus far, we have used the control formula to explain the emergence of both increasing and unimodal PDRs. Density-dependent mortality is a key mechanism for both, but whether an increasing or unimodal pattern emerges depends on the community structure, specifically the strength of apparent competition from a common predator. A structural shift between the two PDRs could thus result from ecological or evolutionary changes that lead to predator extinction, colonization, or increasing predator specialization. This suggests a testable hypothesis: in systems where predation pressure declines (e.g., through predator loss), diversity should shift to increase monotonically with productivity. We are not aware of any experimental study that directly tests this prediction, but this hypothesis could be tested with ecosystem manipulation experiments where predators are added or removed systematically.

More generally, our framework indicates that the strength of apparent competition from shared predation, not solely its presence or absence, has a strong control on the emergent PDR. We next illustrate the transition from the increasing to the unimodal PDR as a function of this control. We simulate ecosystems where species compete for a single limiting resource while varying the number of predator species (Fig. 4a). We assume that each predator targets a distinct group of prey (Fig. S5), with adjacent predators sometimes overlapping in prey species. When the number of predators is small (<10 in our simulations), apparent competition from shared predation is strong, and the unimodal productivity–diversity curve emerges (Fig.4a). As the number of predators increases and their diets become more specialized, apparent competition is reduced, and the curve shifts toward an increasing pattern, eventually converging to the pattern seen in bottom-up guilds where predation is entirely negligible. Thus, species-specific predation (when the number of predators reaches 100 in our simulations) also generates an increasing PDR. Stronger density dependent mortality modulates the quantification of this curve and generally promotes higher species diversity at lower levels of productivity (Fig. S6).

### 4.4 Insights into observed patterns

Finally, we relate our framework and analysis of PDRs to observations. We analyze two plant community datasets that report contrasting PDRs (Fig. 4, (Adler *et al*. 2011; Fraser *et al*. 2015)). We pool all data together across sites, with total biomass used as a proxy for productivity (Supplementary Material). We find that statistically, the data from Adler et al. 2011 supports a significantly increasing PDR (the dashed line in Fig. 4b), whereas the data from Fraser et al. 2015 exhibits a unimodal (hump-shaped) relationship (the dashed line in Fig. 4c). However, despite these differences, the patterns of the two datasets pooled across sites seem visually indistinguishable, highlighting the difficulty of discerning the PDR based on empirical data alone. Additionally, both original studies found that within-site regressions reveal a diversity of patterns, which we also recovered, with increasing (“positive linear” in Fig. 4bc) and unimodal (“concave-down”) patterns as well as decreasing and concave-up patterns, and that many sites indicated no significant relationship.

Our framework provides insight into these variable PDRs (Fig. 4). For one, our simulations help to make it obvious that sampling only part of the productivity gradient can produce diverse PDR patterns (Fig. 4a). For example, a decreasing relationship would be inferred if only the “right-hand-side” component of the unimodal curve were observed. This sub-sampling could explain much of the within-site PDR variability. Overall, our simulations suggest three general patterns. First, unimodal PDRs are most likely to emerge when productivity spans a wide range, because the persistence of a common predator requires a sufficient level of productivity. Resolving the unimodal curve empirically thus requires sampling across a sufficiently broad range, including sufficiently low, intermediate, and high levels of productivity. Second, increasing PDRs are more likely to be observed at low productivity levels in food webs, even with common predators (representing the “left-hand side” of a unimodal curve), as well as across a wide range in productivity in food webs with specialized predators or no predators. Third, declining PDRs tend to occur at relatively high productivity levels, where apparent competition is dominant throughout the entire range. In summary, we posit that density-dependent mortality may be the mechanism underlying many emergent PDRs, with their differences controlled by the strength of apparent competition from shared predation.

## 5. Outlook

We have presented a quantitative framework for understanding how density-dependent mortality shapes community structure and diversity. By extending resource competition theory, we quantified the specifics of how density dependence reshapes the boundary between competitive exclusion and coexistence, enlarging coexistence zones along resource gradients. Moving beyond graphical analyses of a few species, we derived a general control formula for two fundamental types of guilds within complex ecosystems: bottom-up guilds, where species share a resource, and top-down guilds, where species share a predator. This formula predicts species abundance distributions and attributes the abundance of any species to three mechanistic components: the average biomass set by the shared resource or predator, its deviation from the community-averaged net growth rate, and the strength of density-dependent mortality.

Our modular approach, examining tractable subsets (guilds) of complex ecosystems, also has practical implications. Empirically, the net growth rate can be estimated as the coefficient of the linear term in fits of species densities from time-series observations. For example (Eqn. 14), the net growth rate *G*_*i*_ corresponds to the linear term in the following formulation:

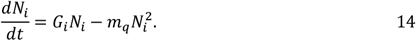

These parameters can in principle be estimated from time-series data without requiring detailed knowledge of specific trophic interactions. This allows our control formula to forecast the population levels of coexisting species using empirical datasets.

We used our control formula to provide a unifying explanation for the emergence of multiple diversity–productivity curves, depending on the food web details. We found that density-dependent mortality is a mechanism that may underly PDRs, and we outlined the dynamics by which this mechanism may result in the emergence of the two most commonly observed forms – increasing and unimodal curves. This obviates the need to rely on environmental heterogeneity and other explanations as mechanisms (though they may also contribute). When predation is negligible or highly specialized, and thus apparent competition is negligible, increasing resource supply allows more species to persist, resulting in a monotonic increase in species richness with productivity. In contrast, when density is additionally constrained by shared predation, generating apparent competition, diversity peaks at intermediate productivity.

More generally, our results contribute to the necessary work of extending theoretical analysis of ecosystem structure to complex ecosystems, such as microbial communities with thousands of species. The control formula provides a tractable framework for predicting species abundances based on traits and measurable quantities, rather than phenomenological interaction coefficients used by the Lotka-Volterra equations. In this way, our mechanistic framework paves the way towards predictive capability in complex ecosystems.

## Acknowledgements

EJZ acknowledges support from NSF (Grant #2125142) and the Simons Foundation Early Career Investigator in Aquatic Microbial Ecology and Evolution Award. LX and EZ acknowledge the Carnegie Institution for Science for funding and support. CAK was supported by National Science Foundation grant EF-2124800.

## Text box 1

In a food web with two species competing for two nutritionally substitutable resources, density-dependent mortality affects the four regions of the resource plane as follows (Fig. 1a-c):

- **Region of no species**: This zone is bounded by the invasion ZNGIs of the two species and two resource axes. In this zone, the supply of both resources is severely limited and cannot support the persistence of either species. This zone is not changed when incorporating density-dependent mortality because the minimum requirement of resources is not impacted: the invasion ZNGIs remain constant as Eqn. 1a-b do not depend on *m*_*q*_ when *N*_*i*_ = 0.
- **Regions of competitive exclusion**: In these regions, resource 1 (or 2) is sufficiently supplied, which allows species 1 (or 2) to maintain its presence. In contrast, the limited supply of resource 2 (or 1) hinders the survival of species 2 (or 1), resulting in its exclusion. The presence of density-dependent mortality narrows this zone and allows species 2 (or 1) to eventually persist with the increase of the supply concentration of resource 2 (or 1).
- **Region of coexistence**: This region is bounded by two boundaries characterized by adequate resource supply levels for both resources. Consequently, it creates an environment where both species can coexist. Density-dependent mortality significantly expands this zone (Fig. 1c).

In a food web with two species *N*_*i*_ competing for one resource *R* and consumed by a common predator *P*, density-dependent mortality affects the four regions of the resource plane as follows (Fig. 1j-l):

- **Region of no species**: When 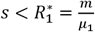, resource supply is insufficient to meet the minimum requirement for either species, even for the best resource exploiter. No species can persist in this range of resource supply rate. Incorporating density-dependent mortality does not change this threshold.
- **Region of species 1 winning**: When 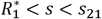, species 1 can persist, but not species 2. Specifically, in the range of 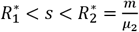, species 2 cannot persist because the resource supply concentration is below its minimum requirement. When 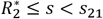, species 2 is excluded by competition with species 1. Density-dependent mortality decreases *s*_21_ (Eqn. 4), which allows species 2 to invade at a lower resource supply compared to the case of no density-dependent mortality.
- **Region of coexistence**: When *s*_21_ < *s* < *s*_12_, both species can coexist. The effect of density-dependent mortality expands the coexistence region through increasing *s*_12_ and decreasing *s*_21_ with increasing *m*_*q*_ (Fig. 1k,l).
- **Region of species 2 winning**: In the last region where *s* > *s*_12_, only species 2 can survive. Increasing resource supply raises the total density of consumer species, resulting in an increase in the density of the shared predator. This apparent competition leads to the exclusion of species 1.

## Text box 2: The control formula derivation and detail

### Derivation overview

There are two ways to derive the density of species at equilibrium. The most direct approach is to solve Eqn. 7 for equilibrium. Specifically, at equilibrium, the density of the consumer can be obtained by setting Eqn. 7a to 0:

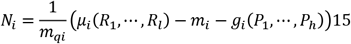

This equation shows that consumer density increases with gains from resources but decreases due to linear mortality and predation. Moreover, density is inversely proportional to the nonlinear mortality rate. While this formulation is broadly applicable in general food webs, it does not reveal how interactions among species within trophic levels contribute to changes in density.

To address this, we instead apply a contrast-based decomposition. In this approach, we derive formulas specifically for the bottom-up and top-down guilds (see Supplementary Material Section 1) and then present a general control expression as the shared functional form. The resulting equation differs from Eqn. 15—particularly in the weighting terms. The advantage of this derivation – our control formula – provides a richer ecological interpretation, as discussed in the main text and summarized in Eqn. 8 and 18.

### The control formula for top-down trophic guilds

Both the bottom-up and top-down guild control formula have the same functional form, the general control formula. In the main text, we provide the specifics for the bottom-up guild, and here we provide the analogues specifics for the tow-down guild. In a top-down trophic guild, all species are preyed upon and largely controlled by a common predator, with unspecified resources that do not constrain total density of consumer species, given by (Fig. 2b):

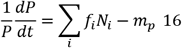

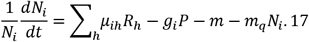

In this case, the explicit form of the control formula Eq. 7 is given by

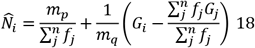

where the average biomass density reflects the control from the predator as 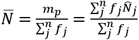, which is the subsistence concentration of the predator, or the mean density 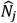weighted by the predation rate *f*_*j*_ . The average community growth rate is weighted by the predator’s assimilation (consumption) rate on each species f as 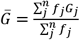, where 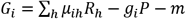. Note that we assume a static (non-oscillatory) equilibrium state, though oscillatory equilibria can be treated by averaging species densities over cycles.

